# Sensitivity to statistical structure facilitates perceptual analysis of complex auditory scenes

**DOI:** 10.1101/126763

**Authors:** Lucie Aman, Samantha Picken, Lefkothea-Vasiliki Andreou, Maria Chait

## Abstract

The notion that sensitivity to the statistical structure of the environment is pivotal to perception has recently garnered considerable attention. Here we investigated this issue in the context of hearing. Building on previous work (Sohoglu & Chait, 2016b), stimuli were artificial ‘sound-scapes’ populated by multiple (up to 14) simultaneous sources (‘auditory objects’) comprised of tone-pip sequences, each with a distinct frequency and pattern of amplitude modulation. Sequences were either temporally regular or random.

We show that listeners’ ability to detect abrupt appearance or disappearance of a source is facilitated when scene sources were characterized by a temporally regular fluctuation pattern. The patterning of the changing source as well as that of the background (non-changing) sources contribute independently to this effect. Remarkably, listeners benefit from regularity even when they are not consciously aware of it. These findings establish that perception of complex acoustic scenes relies on the availability of detailed representations of the regularities automatically extracted from each scene source.

---

Due to their idiosyncratic physical constraints, most animate objects produce statistically structured, temporally predictable, sensory signals (e.g. vocalizations, locomotion). Accumulating evidence from both vision and audition, suggest that observers are sensitive to this patterning and use it to understand, and efficiently interact with, their surroundings (e.g. Rohenkohl et al, 2012; Näätänen et al, 2011; Leaver et al, 2009; Lange, 2013; Andreou et al, 2011; Yaron et al, 2012; Nelken, 2012; Costa-Faidella et al, 2011; Winkler et al, 2009).

Specifically in the context of hearing, It has been proposed that the auditory system’s capacity to rapidly detect predictable patterns in the unfolding sound input may facilitate listening in crowded environments (Winkler et al, 2009; Bendixn et al, 2010; Andreou et al, 2011; Bendixen et al, 2014). We recently provided evidence for this proposal in a magnetoencephalography (MEG) study (Sohoglu & Chait, 2016b), using a stimulus that mirrors the complexity of crowded natural acoustic scenes and an ecologically relevant listening task. Listeners were presented with artificial acoustic scenes (Figure 1) comprising several concurrent sound-streams (‘sources’), each consisting of a sequence of tone pips of a particular frequency and with rates commensurate with those characterizing many natural sounds. Their task was to detect occasional changes (appearance of a source) within those ‘soundscapes’. Keeping the spectral information matched, the temporal structure of the streams comprising the scene was manipulated such they were either temporally regular or random (repeated with a random inter-pip-interval). We demonstrated that activity in auditory cortex rapidly (within 400 ms of scene onset) distinguishes scenes comprised of temporally-regular (REG) vs. temporally-random (RAND) streams. This is manifested by increased sustained activity to REG relative to RAND scenes. Over and above this, appearance of a source in REG scenes is associated with increased responses relative to RAND scenes, mirroring the behavioural data which showed increased accuracy and quicker response times in that condition.

**Figure 1:**
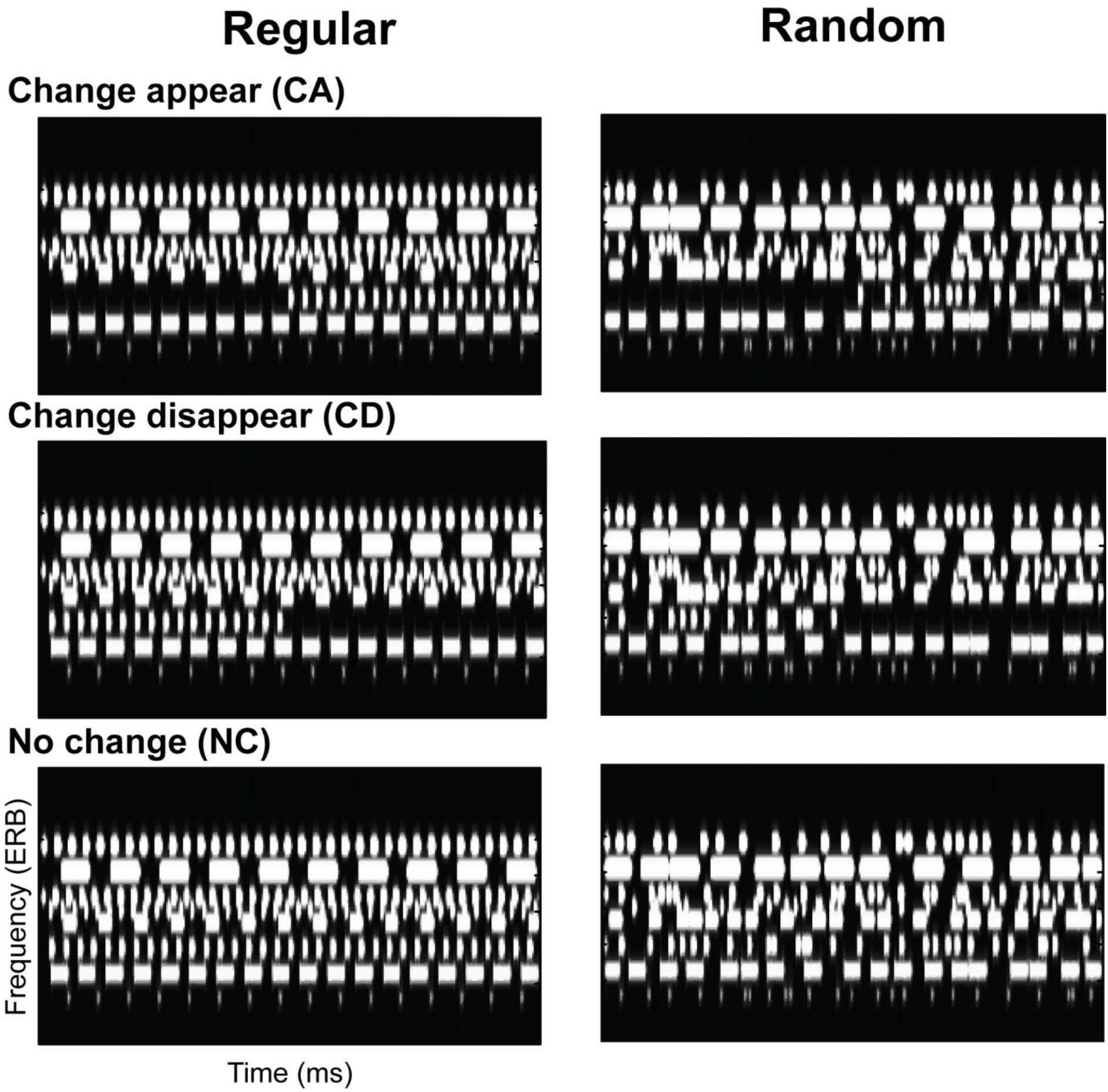
Example of the three variations (‘change appear’, ‘Change disappear’, and ‘no change’) of a scene with 8 sources. Regular (REG) scenes are on the left, random (RAND) scenes on the right. The scenes are matched spectrally, with only temporal structure differing between REG and RAND scenes. The plots represent ‘auditory’ spectrograms, generated with a filter bank of 1/ERB wide channels equally spaced on a scale of ERB-rate. Channels are smoothed to obtain a temporal resolution similar to the Equivalent Rectangular Duration.

The increased sustained activity after scene onset is interpreted as reflecting a mechanism that infers the precision (predictability) of sensory input and uses this information to up-regulate neural processing towards more reliable sensory signals (Feldman and Friston, 2010; Zhao et al., 2013; Auksztulewicz and Friston, 2015; Auksztulewicz et al, 2017; Barascud et al., 2016; Southwell et al, in 2017). According to this ‘precision-weighting’ account, if the auditory system can form precise models about the content of ongoing scenes, novel events that violate those models (e.g. the appearance events studied in Sohoglu et al 2016: see also Southwell et al 2018) would evoke greater neural responses and be perceived as more salient, consistent with the observed change-evoked brain responses and the behavioural advantage associated with REG scenes.

In the series of behavioural experiments reported here, we aim to understand the factors which underlie listeners’ sensitivity to the statistical structure of complex soundscapes. In Experiment 1a, we replicate the behavioural results in Sohoglu & Chait (2016b) and further extend them to item disappearance. In Experiment 1b we show that a similar advantage of regularity persists in scenes where the components are not co-localized but presented from different locations spanning 180° around the listener. In Experiment 2a,b we demonstrate that the effects are not restricted to simple isochronous patterns but extend to more complex forms of temporal regularity. In Experiments 3 and 4 we reveal that the effect of regularity is driven by sensitivity to the temporal regularity of the changing (appearing or disappearing) **source** per se, as well as that of the **context** (the other, non-changing, sources in the scene). Overall the results reveal that listeners routinely extract and track the temporal structure of multiple simultaneous acoustic sources, even in very crowded scenes. Delineating this capacity is crucial towards understanding listening in complex environments and may also help explain failure of scene analysis in certain populations.

## EXPERIMENT 1A – CHANGE DETECTION IS IMPORVED IN REG RELATIVE TO RAND SCENES

Sohoglu & Chait (2016b) previously demonstrated that detecting item appearance in REG scenes is improved relative to RAND scenes. This was interpreted as indicating that listeners extract and track the temporal structure of individual scene components, rendering unexpected events, such as those associated with the appearance of a new stream, more surprising, and hence more salient, in REG relative to RAND scenes.

Here we investigate the same process in the context of item disappearance. If listeners indeed monitor the temporal structure of all components in the scene, they should also be faster and more accurate at detecting item disappearance in REG, relative to RAND, scenes. This is because optimal disappearance detection directly depends on an accurate representation of upcoming tone pips - an ideal observer can detect the cessation of a stream at the moment an expected tone pip fails to arrive. Importantly, since the identity of the disappearing source is a-priori unknown, improved performance in REG scenes would require the ability to simultaneously track the temporal structure of all (or at least a large number of) scene components.

### Materials and Methods

#### Stimuli

The artificial ‘sound-scapes’ used here simulate challenges faced by listeners in natural acoustic scenes, in which many concurrent sound sources, each with a distinctive temporal pattern, are heard simultaneously (Snyder et al, 2012; Eramudugolla et al, 2005). Unlike natural sounds, however, the present stimuli are designed such that sources occupy distinct spectral ranges and hence do not energetically mask each other. This enables us to (1) create the optimal conditions for the streams to be perceived as independent auditory objects (2) measure the effect of growing scene size (number of concurrent sources present) independently of increased inter-source masking. Stimuli (Figure 1) were 2000-4000 ms long artificial ‘scenes’ populated by multiple (4, 8 or 14) streams of pure-tones designed to model sound sources. Each source is characterized by a different carrier frequency (drawn from a pool of fixed values spaced at 2*ERB between 100 and 4846 Hz; Moore & Glasberg, 1983), and is furthermore amplitude modulated (AM) by a square wave – such that the source can be seen as a sequence of tone pips. In a previous series of experiments (Cervantes Constantino et al, 2012), we demonstrated that these stimuli are perceived as a composite ‘sound-scape’ in which individual streams can be perceptually segregated and selectively attended to, and are therefore good models for natural acoustic scenes.

It is important to note that in the present experiments the concept of scene size is confounded with scene density - the more streams (‘objects’) in the scene the closer they are to each other. This is an inevitable consequence of using a fixed frequency range for the scenes, but, importantly, the same constraint also characterizes the notion of scene size in the environment (where the limit may be imposed by the audible hearing range in humans). We continue to refer to the manipulation as ‘scene size’ to be consistent with previous work (Sohoglu & Chait, 2016a; Cervantes Constantino et al, 2012; Gregg & Snyder, 2012; Pavani & Turatto, 2008; Gregg & Samuel, 2008; Eramudugolla et al, 2005). Importantly, the restriction that streams are at least 2 ERB apart controls the issue of density to some extent in that the large spectral separation between neighbouring streams minimizes peripheral masking, enabling the investigation of the effects of increasing scene size without the confound of increasing inter-stream sensory masking.

In the ‘regular’ scenes (REG), the duration of a tone pip (values uniformly distributed between 20 and 160 ms) and the silent interval between pips (values uniformly distributed between 2 and 160 ms) are chosen independently and then fixed for the duration of the scene so that the pattern is regular (see Figure 1, left column). This pattern mimics the regularly modulated temporal properties of many natural sounds. In ‘Random’ (RAND) scenes, tone duration remains fixed throughout the scene, but the silent intervals between successive pips are varied randomly (values uniformly distributed between 2-160 ms) resulting in an irregular pattern (See Figure 1, right column).

Scenes in which each source is active throughout the stimulus are referred to as ‘no change’ stimuli (NC). Additionally, we synthesized scenes in which a source became active (appeared) or inactive (disappeared) at some intermediate time during the scene. These are referred to as ‘change appear’ (CA) and ‘change disappear’ (CD) stimuli, respectively. The timing of change varied randomly (uniformly distributed between 1000 ms and 2000 ms post scene onset), but with the following constraints: The nominal time of change for CA objects coincided with the onset of the first tone while for CD objects the nominal time of change was set to the offset of the last tone augmented by the inter-tone interval, i.e. at the expected onset of the next tone, which is the earliest time at which the disappearance could be detected. For disappearing sources in RAND scenes, it is impossible to define change time in this way (because there is no regular temporal structure). For the purpose of measuring RT, the CD change time in RAND scenes was set to the offset of the last tone-pip augmented by the mean inter-pip-interval (85 ms). Because the distribution of inter-pip-intervals was identical in REG and RAND conditions, if temporal structure does not play a role in change detection, RT should be identical in both conditions.

The set of carrier frequencies and modulation patterns was chosen randomly for each scene, but to enable a controlled comparison between conditions, NC, CA and CD stimuli were generated as triplets sharing the same carrier frequencies and modulation patterns (but differing by the appearance or disappearance of a source; see Fig. 1). They were then presented in random order during the experiment, blocked by change type (NC and CA or NC and CD) and scene type (RAND or REG). Each block contained equal numbers of no change (NC) or change (CA or CD) scenes such that the occurrence of change (and change time) were unpredictable.

Stimuli were synthesized with a sampling rate of 44100 Hz and shaped with a 30 ms raised cosine onset and offset ramp. They were presented with an EDIROL UA-4FX sound card (Roland Corporation) over headphones (Sennheiser HD 555) at a comfortable listening level (~60-70 dB SPL), self-adjusted by each participant. Stimulus presentation was controlled using the *Cogent* software (http://www.vislab.ucl.ac.uk/cogent.php).

#### Procedure

the experiment was conducted in an acoustically-shielded booth (IAC, Winchester, UK). Experimental sessions lasted about 2 hours and consisted of a short practice session with feedback, followed by the main experiment without feedback, divided into runs of approximately 10 minutes each. Subjects were instructed to fixate at a cross presented on the computer screen, and perform a change detection task whereby they pressed a keyboard button as fast as possible when they detected a change in the presented stimulus. They were allowed a short rest between runs.

#### Analysis

Dependent measures are hit rates (number of changes correctly identified), d’ scores and response times (RT; measured between the nominal time of change and the subject’s key press). Since hit rates and d’ reveal consistent effects, in subsequent experiments we report d’ and RT only. The α level was *a priori* set to 0.05.

#### Participants

Ten paid participants took part in the experiment (5 female; mean age = 25 years). All reported normal hearing and no history of neurological or audiological disorder. Experimental procedures (here and in subsequent experiments) were carried out in accordance with the protocols approved by the research ethics committee of University College London, and written informed consent was obtained from each participant. The sample size (10 participants; here and in subsequent experiments) is based on previous experience in the lab with similar data. As can be seen below the effects are stable and consistent across participants, suggesting the sample size is appropriate.

### Results

Figure 2 shows change detection performance for ‘appearing’ (CA; in red) and ‘disappearing’ (CD; in blue) events. For **hit rates** (Fig 2A), a repeated measures ANOVA with scene type (REG vs. RAND), change type (CA vs. CD) and scene size (4, 8, or 12 objects) as factors revealed main effects of scene type (F(1,9)=53.1, p<0.0001), change type (F(1,9)=87.9, p<0.0001) and scene size (F(2,18)=145.3, p<0.0001) as well as the following interactions: scene type × change type (F(1,9)= 9.2, p=0.014; for CD scenes, the difference between REG and RAND was greater than for CA), scene type × scene size (F(2,18)=6.9, p=0.006; for REG scenes, the effect of scene size was smaller than for RAND scenes) and change type × scene size (F(2,18)=28.5, p<0.0001; due to CD showing a steeper decline in performance with scene size than CA).

**Figure 2:**
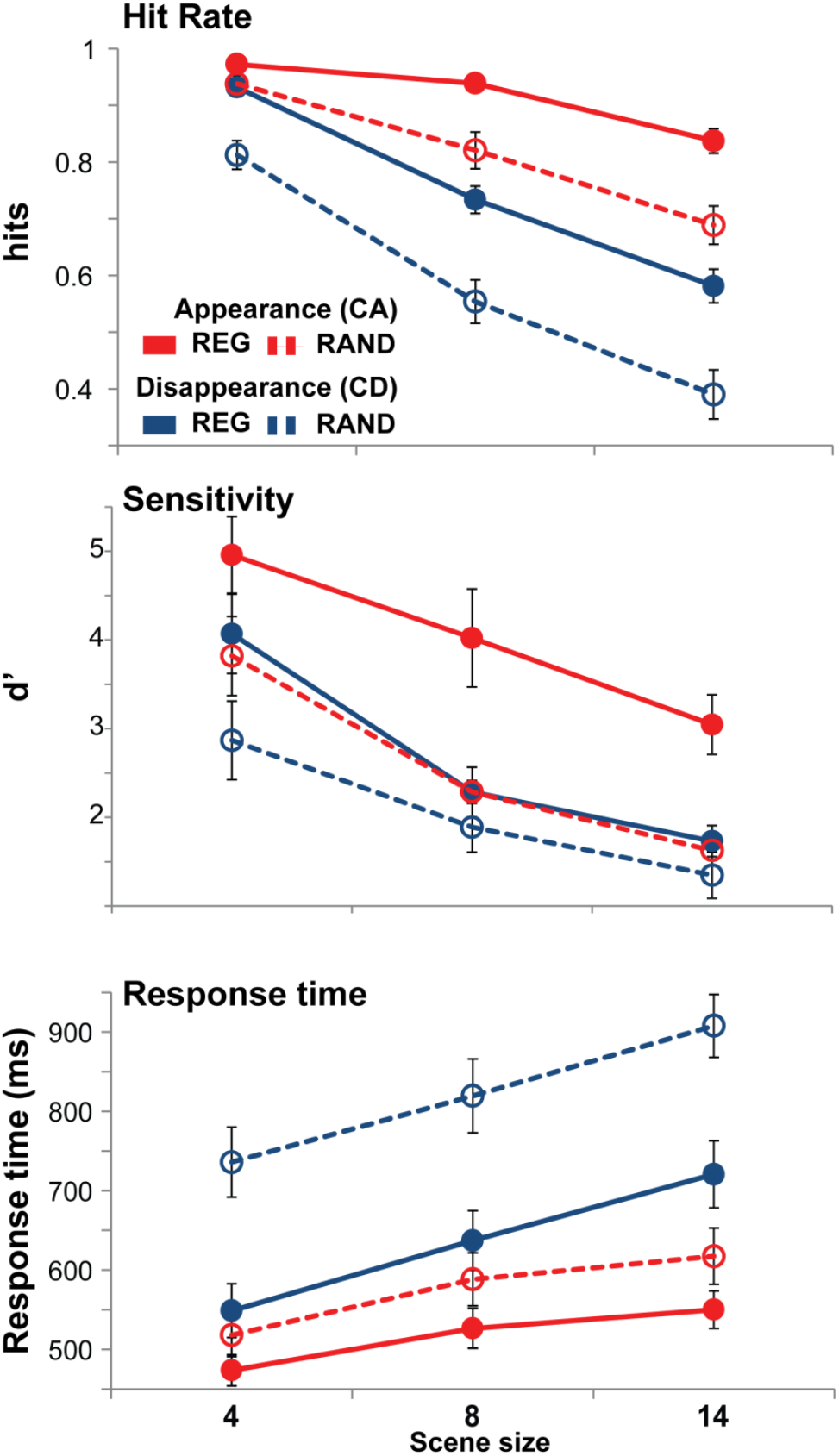
Results of Experiment 1A. Error bars are 1 standard error (SE). In all measures (hit rate, d’ and response time) performance is significantly reduced in RAND relative to REG scenes.

The data are consistent with previous reports that changes associated with appearance of objects in the scene are easier to detect than those associated with disappearances (see discussion in Cervantes Constantino et al, 2012; Sohoglu & Chait 2016a; Pavani & Turatto, 2008). Given the spectral separation between objects, the steep drop in performance for larger scenes is likely associated with the growing computational load of monitoring multiple streams in parallel (rather than inter-component masking). Importantly, the main, novel result is that listeners’ capacity to detect changes (both appearances and disappearances of objects within the scene) depends on temporal regularity.

***d’* sensitivity** scores mirror the hit rate results. A repeated measures ANOVA with scene type, change type and scene size as factors revealed main effects of scene type (F(1,9)=45.3, p<0.0001), change type (F(1,9)=10.8, p=0.009) and scene size (F(2,18)=64.3, p<0.0001) with no interactions. Since d’ an hit rates provide consistent results and Since the d’ measure is a more reliably measure of sensitivity that will be the measure reported in the remaining experiments.

The **response time** data demonstrated a pattern similar to that for the detection performance. Listeners were slower to detect disappearance (relative to appearance) events, and, importantly, for both CA and CD, reaction times were significantly slower in RAND, relative to REG, scenes. A repeated measures ANOVA revealed main effects of scene type (F(1,9)=27.4, p=0.001), change type (F(1,9)=71.5, p<0.0001) and scene size (F(2,18)=85, p<0.0001) as well as the following interactions: scene type × change type (F(1,9)=21, p=0.001; due to performance on CD showing a larger increase in RT than that on CA) and change type × scene size (F(2,18)=10.5, p=0.001).

Thus the results replicate the behavioural results from Sohoglu & Chait (2016b; for CA) and further extend them to demonstrate that CD performance is also improved in REG relative to RAND scenes.

## EXPERIMENT 1B – THE PERCEPTUAL ADVANTAGE OF REGULARITY EXTENDS TO SPATIALIZED SCENES

The results of Experiment 1A demonstrate that listeners rely on temporal regularity in the course of scene analysis. Do they track the regularity of individual sources, or is the observed improvement for REG scenes driven by a representation of a complex temporal regularity across the entire frequency range?

Because regularity of individual components is inherently correlated with overall regularity, this question is difficult to address. However, key elements of the paradigm, including the use of multiple, random-phase streams that are widely spaced in frequency, were explicitly implemented to encourage listeners to process the signals as multiple concurrent ‘auditory objects’. Because the regular patterns characterizing each source are simple, relative to the much more complex aggregate pattern, it is parsimonious to suppose that patterns were extracted within each component separately (e.g. Dau et al 1997a,b). The experimental results, demonstrating very rapid response times in REG scenes provide further support for this assumption.

To further probe this issue, in Experiment 1B we repeated essentially the same paradigm as in Experiment 1A, but in the context of a spatialized scene: Each stream was presented through a different loudspeaker (12 overall; 15° separation) positioned around the listener (See Figure 3). We reasoned that the introduction of spatial separation between individual scene sources will increase their distinctiveness (Bregman, 1990) thereby impeding the representation of any global regularities. If effects of regularity indeed persist in this setting, this would strengthen the conclusion that listeners track the regularity of individual sources.

**Figure 3:**
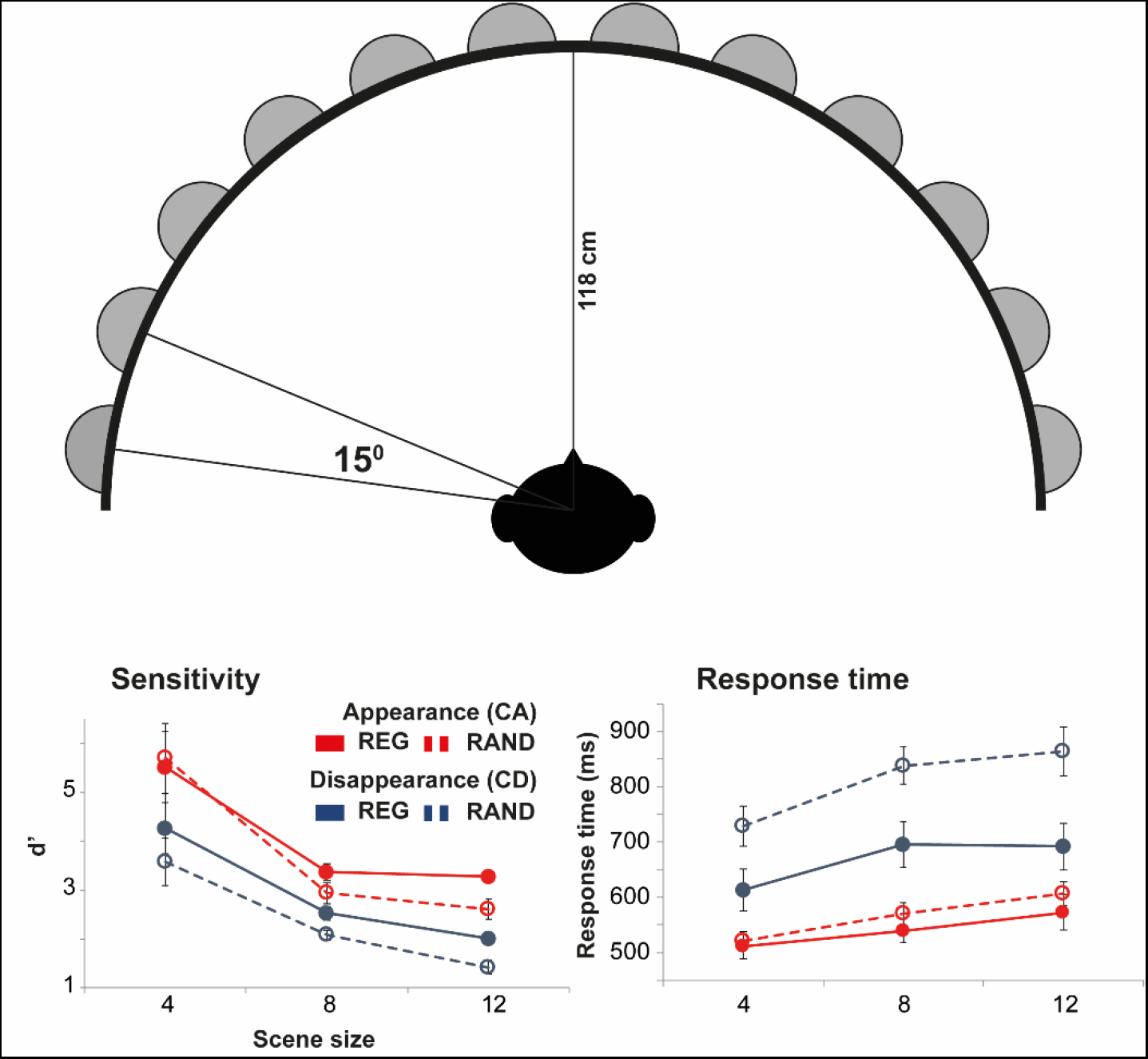
Experiment 1B. [top] Schematic diagram of the loudspeaker array. [bottom] Results of Experiment 1B. Error bars are 1 standard error (SE). In all measures (d’ and response time) performance is significantly reduced in RAND relative to REG scenes.

### Materials Methods & procedure

#### Stimuli

The stimuli were as in Experiment 1A, except for the following differences: The experiment was conducted in an anechoic chamber (IAC, Winchester, UK). Listeners sat in the centre of a 12-loudspeaker array (see Figure 3), with loudspeakers arranged at 15° spacing on the horizontal plain at the level of the listener’s ears. Participants were instructed to fixate at a cross drawn between the two front-most loudspeakers. Scene sizes of 4, 8 or 12 concurrent streams were used.

Scene were generated randomly as before except that, in addition, the location of each stream was randomly assigned to one of the 12 loudspeakers. Therefore, on each trial, the occurrence of a change, the identity of the changing stream. or its location, were unpredictable.

#### Analysis

Dependent measures are d’ scores and response times (The α level was *a priori* set to 0.05.

#### Participants

Ten paid participants took part in the experiment (7 female; mean age = 23.9 years). All reported normal hearing and no history of neurological or audiological disorder.

### Results

Figure 3 shows change detection performance in Experiment 1B. Though it appears that the difference between REG and RAND scenes was smaller in this experiment, a repeated measures ANOVA on d’ data with scene type (REG vs. RAND), change type (CA vs. CD) and scene size (4, 8, or 12 objects) as within subject contrasts and experiment as a between subjects contrast revealed only main effects of scene type (F(1,18)=25.6, p<0.0001), change type (F(1,18)=52.5, p<0.0001) and scene size (F(1,18)=104.9, p<0.0001) with no interactions. This suggest that the introduction of spatial separation between sources did not affect performance. Specifically, REG scenes still yielded better performance than RAND scenes.

Turning to RT: a repeated measures ANOVA on RT data with scene type (REG vs. RAND), change type (CA vs. CD) and scene size (4, 8, or 12 objects) as within subject contrasts, and experiment as a between subjects contrast revealed main effects of scene type (F(1,17)=22.9, p<0.0001), change type (F(1,17)=46.4, p<0.0001) and scene size (F(1,17)=95.1, p<0.0001) with no interactions.

Overall the results of Experiment 1B suggest that the advantage of regularity persists also in scenes where sources are widely spatially distributed.

## EXPERIMENT 2A, B - THE PRECEPTUAL ADVANTAGE OF REGULARITY EXTENDS TO MORE COMPLEX TEMPORAL PATTERNS

Experiment 1 used simple isochronous patterns. Here we ask whether an advantage of regularity will also extend to more intricate patterns. Towards this aim we created non isochronous, repeating patterns (Figure 4) which model increasingly complex temporal regularities.

**Figure 4:**
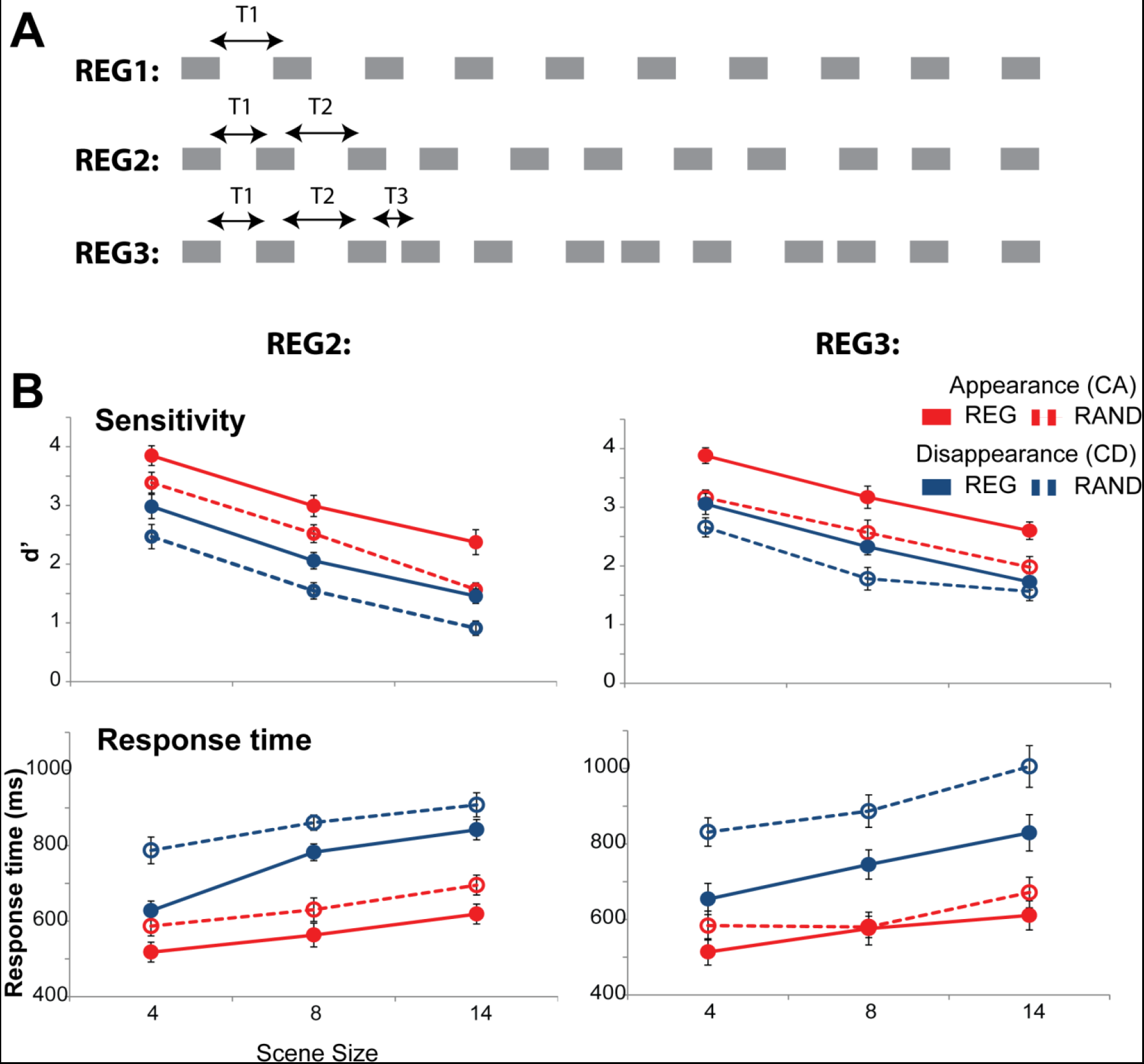
Experiments 2a and 2b. A Schematic representations of the regular patterns used. Scene sources in Experiment 1 (REG1) contained a fixed inter-tone-interval (T1) that was randomly chosen for each source in each trial. Those in Experiment 2a (REG2) contained two different, regularly repeating, inter-tone-intervals (T1 and T2). T1 and T2 were randomly chosen for each source in each trial. REG patterns in Experiment 2b (REG3) contained three different, regularly repeating, inter-tone-intervals (T1, T2 and T3). These were randomly chosen for each source in each trial. B Results of Experiment 3a (Left) and Experiment 3b (Right) expressed in terms of d’ scores (top) and reaction times (bottom). Error bars are 1 standard error (SE). As in Experiment 1, performance is significantly increased in REG relative to RAND scenes.

### Materials and Methods

#### Stimuli

Stimuli were identical to those in Experiment 1, above, except that REG scenes were characterized by increasingly complex temporal patterns. In **Experiment 2A** REG sources were constructed by randomly choosing two inter-tone-interval durations (T1, and T2 in Figure 4A; values uniformly distributed between 20 and 160 ms) which then repeated regularly. Henceforth, this condition will be referred to as REG2. In **Experiment 2B** scene sources contained three randomly selected, regularly repeating inter-tone-intervals. This condition is referred to as REG3. In both experiments, only inter-tone intervals were manipulated. Tone-pip durations were randomly chosen for each source, from within the same range as above, and then fixed for the duration of the stimulus. Inter-tone-interval durations and tone-pip durations were chosen anew for each source in each trial.

#### Participants

Eleven subjects (mean age 22.9 years; 7 females) participated in Experiment 2A and 10 subjects (mean age 22.1; 9 females) participated in Experiment 2B. An additional participant was excluded from the analysis due to very low performance scores (d’=1.5 in the easiest condition [REG CA scene size 4]; for the rest of the subjects d’>3.3). Four subjects participated in both experiments.

### Results

Results are presented in Figure 4. Performance in both experiments resembled that in Experiment 1. REG2: a repeated measures ANOVA on **d’** with scene type (REG vs. RAND), change type (CA vs. CD) and scene size (4, 8, or 14 objects) as factors revealed main effects of scene type (F(1,10)=32.3 p<0.0001), change type (F(1,10)=92.6 p<0.0001) and scene size (F(2,20)=166.5 p<0.0001) with no interactions. An identical pattern is observed for REG3: main effects of scene type (F(1,9)=16.6 p=0.003), change type (F(1,9)=79.6 p<0.0001) and scene size (F(2,18)=82.9 p<0.0001) with no interactions.

**Response time** data revealed effects comparable to those in Experiment 1. REG2: a repeated measures ANOVA on RT data revealed main effects of scene type (F(1,10)=28.87 p<0.0001), change type (F(1,10)=88.1 p<0.0001) and scene size (F(2,20)=39.67 p<0.0001) with no interactions. REG3: main effects of scene type (F(1,9)=85 p<0.0001), change type (F(1,9)=351.6 p<0.0001) and scene size (F(2,18)=32.5 p<0.0001) as well as the following interactions: change type x scene size (F(2,18)=11 p=0.002; due to CD showing larger decline in performance with scene size than CA) and scene type x change type (F(1,9)=17.3 p=0.002). Post hoc tests revealed significant effects of scene type for both CA and CD (F(1,9)=11.8 p=0.007; F(1,9)=54 p<0.0001), suggesting the interaction is due to differing marginal means.

An across group ANOVA was also conducted to compare performance for REG2 and REG3. For d’ this revealed main effects of scene type (F(1,19)=46.037 p<0.0001), change type (F(1,19)=168.8 p<0.0001) and scene size (F(2,38)=239 p<0.0001) and an interaction of group (Exp2A vs Exp2B) x Scene size (F(2,38)=4.2 p=0.024. The interaction is due to a smaller scene size effect in the Exp2B group, which is likely attributable to individual differences. A similar test for RT demonstrated main effects of scene type (F(1,19)=91 p<0.0001), change type (F(1,19)=295.6 p<0.0001) and scene size (F(2,38)=70.9 p<0.0001) as well as the following interactions: change type x scene size (F(2,38)=10.4 p<0.0001); scene type x change type (F(1,9)=18.58 p<0.0001); scene type x change type x group (F(1,19)=6.61 p=0.019). A similar analysis on false positive rates (data not shown) showed main effects of scene type (RAND>REG; F(1,19)=8,19 p=0.07), scene size (F(2,38)=5.17 p=0.01), as well as the following interactions: scene type x scene size (F(2,38)=8.586 p=0.01); change type x scene size (F(2,38)=8.96 p=0.001).

Overall, the data demonstrate that, on all measures, change detection performance was significantly improved in REG relative to RAND scenes. This suggests that listeners are able to track complex regular patterns associated with multiple simultaneous acoustic sources.

## EXPERIMENT 3 – SOURCE- AND CONTEXT- REGULARITY CONTRIBUTE INDEPENDENTLY TO IMPROVED PERFORMANCE

Next, we sought to determine whether the effect of regularity is driven by sensitivity to the temporal regularity of the changing (appearing or disappearing) **source** per se, or by that of the **context** (the other, non-changing, scene elements). We reasoned earlier that sensitivity to context regularity is key for detecting appearance events (CA). This is because the high predictability of the unfolding REG scene renders the onset of new, unexpected, streams particularly salient. However, since CA detection can, in principle, be based on the first transient associated with the onset of the new stream, it may be that source patterning as such does not affect performance. In contrast, we expect that for item disappearance (CD), sensitivity to source regularity should be of key importance because, as discussed above, detection of the cessation of a stream can be vastly improved by tracking its temporal structure.

To understand the effects of source and context we systematically de-coupled the two factors by creating scenes in which the regularity of the changing component is independent of the regularity of the rest of the scene.

### Martials and methods

#### Stimuli

‘Regular’ (REG) and ‘Random’ (RAND) scenes were created as before (Experiment 1) with the exception that the regularity of the changing (appearing or disappearing) source was manipulated independently of the regularity of the rest of the sources in the scene, resulting in 4 configurations for each change type (CA, CD or NC): REG_REG, REG_RAND, RAND_REG and RAND_RAND (in each case the first term refers to the regularity of the scene context, the second to the regularity status of the changing source). Two scene sizes – 8 and 14 – were used. Stimuli were blocked by context and change type (CA vs CD).

#### Participants

Ten participants (6 female; mean age = 29.4 years) took part in the experiment.

### Results

Results are in Figure 5. A repeated measures ANOVA on **d’** data, with context regularity (REG vs. RAND), source regularity (REG vs. RAND), change type (CA vs. CD) and scene size (8 or 14 objects) as factors, showed main effects of context regularity (F(1,9)=92.8 p<0.0001), source regularity (F(1,9)=21.2 p=0.001), change type (F(1,9)=66.8 p<0.0001) and scene size (F(1,9)=87.4 p<0.0001) as well as the following interactions: change type x scene size (F(1,9)=6.28 p=0.033; also seen in Experiment 1) and context regularity x change type (F(1,9)=25.6 p=0.001). To understand this interaction, separate ANOVAs were run for CA and CD. For CA, this revealed main effects of context regularity (F(1,9)=122.6 p<0.0001) and scene size (F(1,9)=44.2 p<0.0001) but no effect of source regularity (F(1,9)=1.8 p=0.205). In contrast, for CD all three main effects were significant: F(1,9)=31.5 p<0.0001, F(1,9)=47.6 p<0.0001 and F(1,9)=35.3 p<0.0001 respectively.

**Figure 5:**
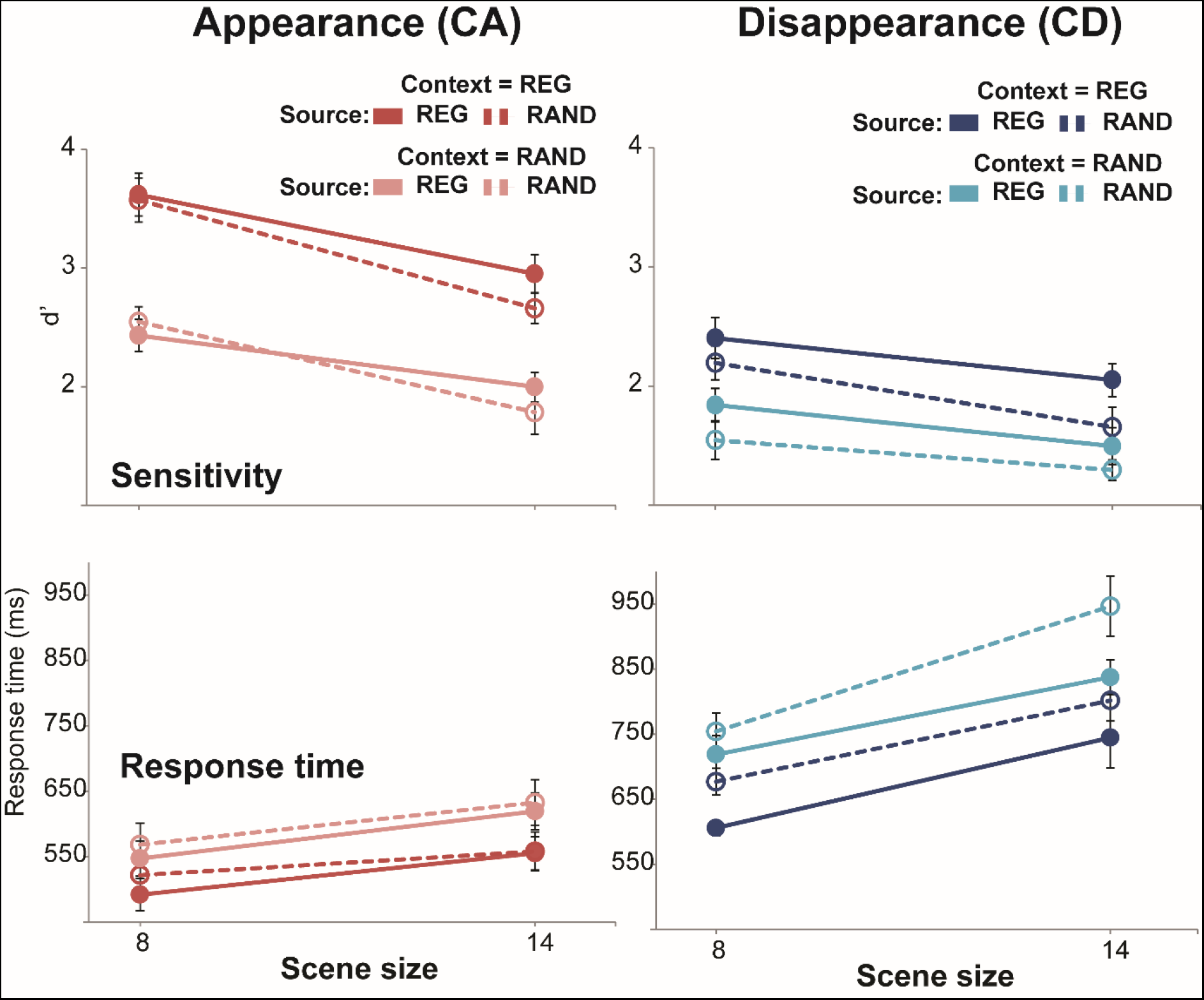
Results of Experiment 3. Appearance changes (CA; red colours) are on the left and disappearance changes (CD; blue colours) are on the right. REG context conditions are in darker colours; RAND context conditions are in lighter colours. REG source conditions are plotted with solid lines; RAND source conditions are plotted with dashed lines. Error bars are 1 standard error (SE). The results demonstrate that the patterning of the changing source as well as that of the background (non-changing) sources both contribute independently to the advantage of regularity in the context of change detection.

Therefore, the data demonstrate that whilst context regularity influenced both CA and CD, source regularity only affected CD performance. This is consistent with the hypothesis that the detectability of CA depends primarily on the first event within the appearing stream (and as such shouldn’t be affected by the patterning of the sequence) whereas successful coding of temporal structure is critical for the rapid detection of source disappearance. Indeed, to efficiently determine that a source has disappeared from the scene, an ideal observer must ‘acquire’ the pattern of onsets and offsets associated with that channel, and respond as soon as an expected tone pip fails to arrive. Importantly, since the identity of the changing source varied randomly from trial to trial, to achieve optimal performance one must be able to represent the temporal structure of *all objects* within the scene. That listeners were indeed consistently better at detecting CD events in REG scenes demonstrates that, listeners do, at least to some extent, acquire the temporal structure of all on-going scene elements and use this information during scene perception.

For **response times**, a repeated measures ANOVA showed main effects of context regularity (F(1,9)=68.4 p<0.0001), source regularity (F(1,9)=26.1 p=0.001), change type (F(1,9)=98.4 p<0.0001) and scene size (F(1,9)=53.6 p<0.0001) as well as the following interactions: change type x scene size (F(1,9)=16.3 p=0.003; also seen in Experiment 1, above, and due to CD showing larger decline in performance with scene size than CA) and context regularity x change type (F(1,9)=10.8 p=0.009). To understand the latter interaction, separate ANOVAs were run for CA and CD. In both cases all three main effects were significant, with no interactions. CA: main effect of context regularity: F(1,9)=17.9 p=0.002, main effect of source regularity: F(1,9)=11 p=0.009, main effect of scene size: F(1,9)=28.9 p<0.0001. CD: F(1,9)=23.7 p=0.001, F(1,9)=19.8 p=0.002, and F(1,9)=42.7 p<0.0001, respectively.

Therefore, RT as a measure of performance suggests that both CA and CD detection are affected by the regularity of the context as well as that of the changing component itself. The source regularity effect for CA, which is observed in RT but not d’, suggests that whilst component regularity does not affect CA detectability per se, it does speed-up its detection.

## EXPERIMENT 4 – REG STREAMS IN A RAND CONTEXT DO NOT POP OUT

The design of experiment 3 was possibly confounded in the sense that regular sources in a random context might have perceptually stood out, even before the actual change event, thus facilitating the scanning for possible changes.

Here we investigated the extent to which listeners are sensitive to such situations: do REG streams in a RAND context (or vice versa - RAND streams in REG scenes) pop out?

### Materials and methods

#### Stimuli

This experiment used only NC stimuli (same length and other parameters as described above). ‘REG context’ scenes contained all regular streams (‘Foil scenes’) or one random stream among regular streams (‘target’ scenes’; 50%). Conversely, ‘RAND context’ scenes contained all random streams or (in 50% of the signals) one regular stream among random streams (‘target’ scenes). Within each context condition, scenes were generated in target/foil pairs such that each duo consisted of identical sources (in terms of frequency and temporal properties) only differing by the temporal structure of the target stream (See Figure 6A). The stimuli were then presented to the listeners in random order, blocked by context type (REG or RAND). Three scene sizes (4, 8 and 14 sources) were used. Participants were instructed to detect the odd sources (regular among random or vice versa – ‘target’ scenes). The task was first explained using a visual depiction of the stimuli (similar to Figure 6 here) and short practice session with feedback (presented on the screen after each trial) was also provided. Unlike the other experiments reported here, feedback was also provided during the main session, to make sure participants are able to perform as well as possible. This was deemed necessary because pilot runs suggested performance is expected to be very low on some of the conditions. Participants were instructed to respond at any time during the stimulus or inter stimulus interval. RT is therefore not analysed here.

**Figure 6:**
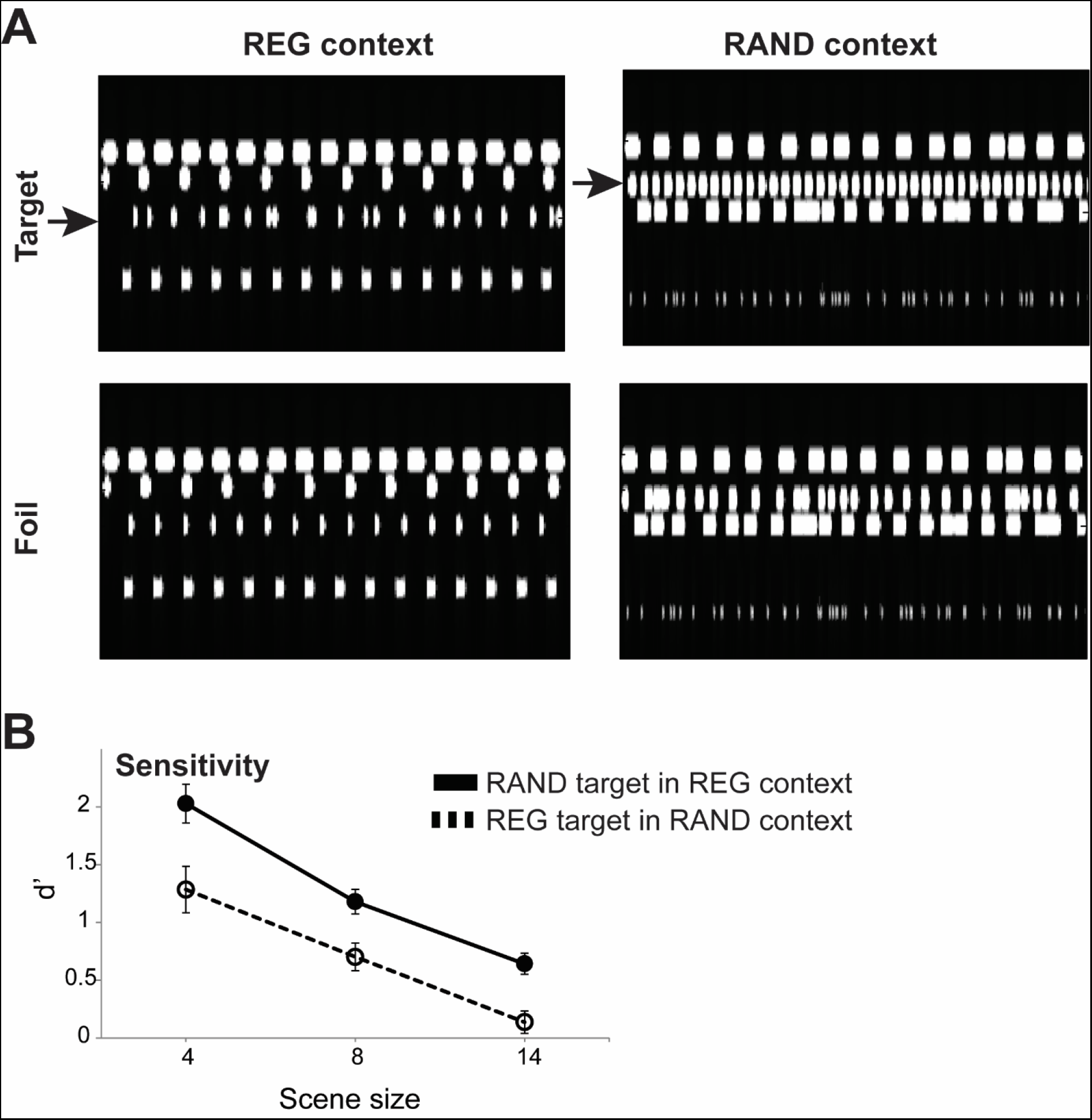
Experiment 4. An Example of REG and RAND context scenes (left and right, respectively) with 4 sources. ‘Foil’ scenes (bottom) contain all REG or all RAND sources; ‘Target’ scenes (top) contain an odd source – regular among random or vice versa, indicated with arrows. The plots represent ‘auditory’ spectrograms, generated with a filter bank of 1/ERB wide channels equally spaced on a scale of ERB-rate. Channels are smoothed to obtain a temporal resolution similar to the Equivalent Rectangular Duration. B Results of Experiment 4. The REG context condition is plotted with a solid line, the RAND context condition is plotted with a dashed line. The results demonstrate that it is consistently easier to detect a random stream among regular streams (REG context) than vice versa.

#### Participants

Nine subjects participated in the experiment (4 female; mean age = 28.7 years). The data from an additional participant were not useable due to a technical error.

### Results

Figure 6B shows the results of Experiment 4. Hit rate and d’ results were qualitatively identical and hence only d’ data are presented. A repeated measures ANOVA on **d’** scores, with context (REG versus RAND) and scene size (4, 8 or 14 objects) as factors, revealed main effects of context (F(1,8)=37.8, p<0.001) and scene size (F(2,16)=52.3, p<0.001); no interactions. No significant effects were observed in the false positive data.

The relatively steep decline with scene size suggests that regularity does not strictly ‘pop-out’ but is rather discovered via some search-based process. That it is overall easier to find a random source in a REG context, rather than a regular source in a RAND context, is in line with the account proposed above: listeners acquire the temporal patterning associated with each source in a REG context scene such that events that do not conform with these patterns (associated with the presence of a RAND stream) are relatively easy to detect. Conversely, in a RAND context, where most events are unpredictable, it is more difficult to spot the one stream that follows a regular pattern.

Importantly, the results demonstrate that at the largest scene size (14) participants are at floor for detecting a regular item in a RAND context (one sample T test against 0: t=1.4, p=0.19). Combined with the findings from Experiment 3, above, the data demonstrate that despite not being aware of the regularity of the sources, participants implicitly used this information for change detection: even with 14 concurrent sources in the scene, subjects’ change detection performance benefitted when a disappearing stream was regular, relative to when it was random. Indeed, all the results of Experiment 3, both in terms of d’ and RT, remain significant when running the ANOVA on scene size 14 only (p<0.01 for all). This reveals that even in very crowded scenes, participants are able to utilize temporal regularity, of which they are not consciously aware, to efficiently detect change events.

## GENERAL DISCUSSION

The auditory system is tuned to changes in the acoustic environment (Cervantes Constantino et al, 2012; Demany et al, 2010). We therefore chose a change detection task as a neuro-ethologically relevant means by which the role of sensitivity to temporal structure in the course of auditory scene analysis can be studied.

We demonstrate that listeners’ ability to detect changes, manifested as the abrupt appearance or disappearance of a source, is facilitated when scene sources are characterized by a temporally regular fluctuation pattern (experiment 1). The patterning of the changing source as well as that of the background (non-changing) sources both contribute independently to this effect (Experiment 3). The advantage of regularity, relative to random temporal patterning, is observed even when using complex, non-isochronous, temporal patterns (Experiment 2). These findings establish that perception of complex acoustic scenes relies on the availability of detailed representations of the regularities automatically extracted from each scene source.

### Are listeners coding local (frequency specific) or global temporal regularities?

An important point concerns whether the patterns were detected within each frequency channel separately, or identified as a complex, temporal regularity across the entire frequency range. In the present stimuli, local regularity is intrinsically linked to global regularity and it is therefore difficult to entirely dissociate the two. Because the regular patterns characterizing each source are simple, compared with the much more complex aggregate pattern, it may be reasonable to conclude that patterns were extracted within each component separately. Several key observations support this assertion: (a) The advantage of regularity persisted even for complex temporal patterns (Experiment 2). (b) The advantage of regularity persisted in the face of spatial separation between streams (Experiment 1B). (c) The ‘source’ effect in Experiment 3 - listeners exhibited improved performance when the changing source was regular even when the rest of the components in the scene were random. Conversely, they showed reduced performance when the changing component was random in an otherwise regular scene.

Mounting evidence suggests that listeners are tuned to the temporal structure of sound sequences and use this information to anticipate and improve their interaction with expected events, even in the absence of directed attention (Barnes & Jones, 2000; Geiser et al, 2012; Andreou et al, 2015). Accumulating work using the ‘omission MMN’ paradigm, demonstrates that the neural machinery is ‘pre-activated’ to process predicted signals (e.g. Hughes et al, 2001; Bendixen et al, 2009; Janata et al, 2001). Importantly, we demonstrate that the auditory system’s ability to track the temporal structure of on-going sound input and register when it is violated persists even when the scene is heavily populated with concurrent objects and the identity of the changing component is in advance unknown.

### Do regular patterns attract attention?

It has been suggested that attention can be understood as a process that infers the level of predictability of sensory signals such that highly predictable sensory streams capture attention in a bottom-up manner (Jones et al, 2002; Arnal & Giraud, 2012; Feldman & Friston, 2010; Schroger et al, 2015). In contrast, here (Experiment 4) we show that regular patterns in a background of random patterns do not pop-out and are in fact always harder to detect than vice versa. This suggests that perception is not drawn to predictability per se but rather to violation of predictability (see also Southwell et al, 2017; Meijs et al, 2018):

Consistent with this, we demonstrate that listeners benefitted from regularity despite not being consciously aware of it: Listeners were at floor when asked to determine whether a regular source was present in a scene containing 14 concurrent random streams (Experiment 4), but exhibited a sizeable improvement to change detection performance when that source disappeared (Experiment 3). This suggests that the temporal structure of that source was automatically tracked by the auditory system and used to facilitate scene analysis. This finding is in line with a previous demonstration by Zhao et al (2013) in the visual modality: A visual search task was facilitated at a location which previously contained a regularity. This occurred even though participants reported not being aware of the regular pattern.

On the whole, the results suggest that while sensitivity to regularity plays a key role in shaping our perception of our surroundings, this does not translate to explicit attentional capture as proposed by some formulations of ‘predictive coding’ (e.g. Feldman & Friston, 2010; see also Southwell et al, 2017). Instead what captures attention are violations of regularity.

### Sensitivity to regularity in the service of auditory scene analysis

In an MEG study (Sohoglu & Chait, 2016b) we recorded responses to REG and RAND scenes (as in Experiment 1, here) in the context of an appearance (CA) detection task. The behavioural advantage associated with REG scenes was accompanied by increased responses in auditory cortex and parietal cortex both before, as well as after, the change. This was interpreted as reflecting the operation of mechanisms which rapidly infer the precision (predictability) of sensory input and upregulate responses to reliable sensory information, such that violations of these patterns (e.g, in the form of an appearing or disappearing sources) evoke higher prediction errors (see also Southwell & Chait, 2018).

The behavioural effects observed here support this interpretation: The ‘context’ effects shown in Experiment 3 (where listeners were better at detecting changes in scenes where the ‘background, non- changing, sources were regular) and the demonstration that listeners are consistently better at spotting random sequences within scenes that otherwise comprised of regular components, than vice versa (Experiment 4) demonstrate increased sensitivity to deviants in REG than RAND context.

In all, these results demonstrate that the auditory system rapidly discovers regular structure in the unfolding sensory input, in line with a broader theoretical framework which views the brain as a regularity extractor (Friston, 2005; Hohwy, 2013; Winkler & Schroger, 2015; Winkler et al, 2009; Zhao et al, 2013; Wacgone et al, 2011; Barascud et al, 2016). Whilst regular structure per se, does not attract attention it is monitored automatically and used to facilitate listening. Transients (onsets of individual tones) in regular streams are predictable and thus do not require substantial processing resources, making it easier to ignore regular patterns when these are task irrelevant (Andreou et al, 2011; Southwell et al, 2017). On the other hand, unexpected events in an otherwise predictable stream are rendered as ‘surprising’ or ‘salient’ and capture bottom-up attention (Kaya & Elhilal, 2014; Huang & Elhilali, 2017; Southwell & Chait, 2018).

It is remarkable that listeners in the present experiments exhibited the ability to track the regularity of sources even in very crowded scenes, populated by many simultaneous sound streams. That normal, young, listeners rely on this capacity so routinely, makes it an interesting feature to investigate in certain clinical populations typically linked to failure to extract temporal regularities (Schapiro et al, 2014; Davalos et al, 2018; Ciullo et al, 2018) as well as during healthy aging (Rimmele et al, 2012).

## Author contribution statement

MC designed the experiments; LA, SP and LVA conducted and analysed the experiments; All authors wrote the manuscript.

## Acknowledgements

This study was supported by a BBSRC project grant to MC.

